# Carbohydrate activity regulation of floral quantity during the juvenile phase in black pepper (*Piper nigrum* L.)

**DOI:** 10.1101/414938

**Authors:** Chao Zu, Jianfeng Yang, Zhigang Li, Can Wang, Huan Yu

## Abstract

Black pepper plants maintain synchronous vegetative growth and flowering during their lifetimes, even in the juvenile phase. How to limit the duration of flowering, facilitate a return to vegetative development should be studied. Light intensity has been reported to affect the levels of stored carbohydrates in some horticultural trees. However, it is unclear whether increased flowering intensity in adaptable light intensity is due to the regulation of carbohydrates in black pepper. Here, we report the characterization of carbohydrates and enzyme turnover in major source leaves under shading treatments during the juvenile phase of pepper. In addition to the previous finding that carbohydrate levels are correlated with flowering time, we report the novel finding that carbohydrate contents in the leaves control floral quantity. To gain insights into the underlying physiological mechanisms, we analyzed the effect of shading on sugar contents and floral transition, which revealed that shading regulated carbohydrate levels, an increase in starch accumulation improved floral quantity, and sucrose-starch ratio played a negative role in inflorescence quantity. Based on this analysis, we characterized the changes in enzyme activities in the leaves that affect carbohydrate dynamics and identified the key indicator enzymes and optimal shading intensity for the five developmental periods.

**Highlight:** We identified the key carbohydrates, indicator enzymes and optimal shading intensity to control inflorescence quantity during the juvenile phase in black pepper for labor-saving.

## Materials and Methods

### Plant materials

This experiment was conducted in the pepper field at the Spice and Beverage Research Institute (SBRI) in the Xinglong region in southeastern Hainan of China (18°72′-18°76′ N, 110°19′-110°22′ E) during 2012-2014. This region is characterized by a tropical monsoon climate (Zu et al., 2016). In this study, *P. nigrum* cv. Reyin-1 in the juvenile phase was used as the material. Artificial shading was imposed during an annual growing period, including the recovery stage (S1), full-bloom stage (S2), expanding stage (S3), grain-filling stage (S4) and fruit maturation stage (S5), with one to four black polyethylene shading nets placed 50 cm above the pepper canopy, resulting in shading intensities of 15%, 30%, 60% and 75%, respectively (Zu et al., 2016). No net covering was used as the control (CK). There were three replicates in each shading treatment and 11 trees in each replicate. The peppers were planted in a north-south row orientation with an intrarow distance of 2.5 m and an intratree distance of 2.0 m in 2012. For each replicate, three uniformly grown pepper trees were selected from which to collect and record data on flowering spikelet quantity every 30 days. In addition, the major source leaves (the second leaf away from the inflorescence) were sampled once in the middle of each stage and were then immediately wrapped in aluminum foil, frozen in liquid nitrogen, placed into a sealed plastic bag, and stored at −80°C. Another three uniformly grown pepper trees were used to collect and determine the vegetative tissue biomass of black pepper. The others were backup trees.

### Leaf carbohydrate content determination Soluble sugar and sucrose extraction and analysis

A total of 0.25 g of major source leaves was minced in a centrifuge tube with 0.10 g of activated carbon as an adsorbent. Then, 4 mL of 80% alcohol was added, followed by oscillation at 80°C for 30 min and then cooling and centrifugation. The supernatant was transferred into a test tube with a screw lid. A total of 4 mL of 80% alcohol was added to the sediment, the mixture was centrifuged again, and the supernatant was transferred into the test tube; this step was repeated twice. The alcohol was removed from the supernatant with a centrifugation evaporator, deionized water was added, and the final supernatant after centrifugation was the sample for determination of soluble sugar and sucrose contents. Next, 5 mL of anthrone-sulfuric acid reagent was added to the 1 mL of filtrate and boiled at 100°C for 5 min. Once cooled, the soluble sugar content of the reaction mixture was determined by measuring the OD value at 620 nm. A total of 200 μL of 2 mol/L sodium hydroxide was added to the 1 mL of filtrate and boiled at 100°C for 10 min. Then, 3.5 mL of 30% hydrochloric acid and 0.8 mL of 0.1% resorcinol was added, and the reaction mixture was boiled at 80°C for 10 min. Once cooled, the sucrose content of the reaction mixture was determined by measuring the OD value at 480 nm (Wang et al., 2015).

### Leaf starch extraction and analysis

The alcohol and water were removed from the soluble sugar sediment with a centrifugation evaporator. Then, deionized water was added, the mixture was incubated at 100°C for 15 min and then cooled, 2 mL of 9.2 mol/L perchloric acid was added to the sediment and extracted for 15 min, the sediment was filtered, and the volume of filtrate was adjusted to 25 mL. A total of 2 mL of 4.6 mol/L perchloric acid was then added to the sediment and extracted for 10 min. Next, the filtrate and 6 mL of deionized water were mixed together in a 25 mL volumetric flask. To this 1 mL mixture, 5 mL of anthrone-sulfuric acid reagent was added, and the mixture was immediately boiled at 100°C for 5 min. Once cooled, the starch content of the reaction mixture was determined by measuring the OD value at 620 nm (Wang et al., 2015).

### Leaf carbohydrate metabolism enzyme activity assay Leaf carbohydrate metabolism enzyme extraction

A total of 0.5 g of frozen leaf blade samples was homogenized using a chilled mortar with 0.08 g of PVPP and 4 mL of extraction buffer (50 mM HEPES-NaOH with a pH of 7.5, 20 mM MgCl2, 2 mM EDTA, 0.05% Triton X-100, 2.5 mM DTT, and 1 mM PMSF). The homogenate was centrifuged at 10,000 r/min for 20 min at 4°C.

### SPS and SS activity analysis

For the SPS activity assay, 100 μL of enzyme extract was mixed with 150 μL of extraction buffer (50 mM HEPES-NaOH, 10 mM MgCl_2_, 3 mM UDPG, and 10 mM F-6-P, with a final pH of 7.4). The reaction was initiated by incubating the enzyme extract at 30°C for 30 min. The reaction was stopped by adding 50 μL of 2 M NaOH and heating the solution for 10 min at 100°C to destroy unreacted hexoses and hexose phosphates. Then, the solution was cooled and mixed with 875 μL of 30% (w/v) HCl and 250 μL of 0.1% (w/v) resorcin before being incubated for 10 min at 80°C. Sucrose content was calculated from a standard curve measured at A480 nm. For the SS activity assay, F-6-P in the extraction buffer was replaced by fructose, and the other reaction steps were the same as those in the SPS activity analysis (Liu et al., 2013).

### AI and NI activity analysis

The acid invertase (AI) and neutral invertase (NI) activities were assayed using the method of Batta and Singh (1986), with some modifications. The reaction mixtures consisted of 20 μL of enzyme extract and 180 μL of buffer [0.1 M sodium acetate buffer (pH of 4.8) and 0.1 M sucrose for AI; 0.1 M potassium phosphate buffer (pH of 7.2) and 0.1 M sucrose for NI]. After the mixture was incubated at 37°C for 30 min, 200 μL of DNS was added, and the mixture was boiled for 5 min. The absorbance of the reaction solution was determined at 540 nm (Huang et al., 2013).

### Statistical analysis

All data were the means of at least three replicates with standard errors of the mean. Histograms were developed using GraphPad. Prism. v5.0 (Cabit Co., USA). Variance (ANOVA) was measured using the SAS statistical analysis package (version 8.2, SAS Institute, Cary, NC, USA).

## Introduction

Perennial plants live for many years and repeatedly cycle between vegetative and reproductive development. How perennials undergo repeated cycles of vegetative growth and flowering during the growing periods has not been extensively studied (Wang et al., 2009). Flowering is best understood in annual plants, but the cultivars of *Piper nigrum* L., such as *P. nigrum* cv. Reyin-1, are perennial. They are polycarpic and maintain synchronous vegetative growth and flowering during their lifetimes. During the juvenile phase, flowering can compete with the vegetative growth of pepper vines for nutrients, and the traditional method of picking inflorescences demands a substantial amount of manual labor (Zu et al., 2016). Therefore, how to limit the duration of flowering, facilitate a return to vegetative development, and prevent some branches from undergoing the floral transition should be studied.

Floral initiation is best understood in herbaceous species. Therefore, a discussion of the control of floral initiation in *Arabidopsis* precedes such discussions in trees and vines. Four major flowering pathways have been characterized in herbaceous species, including those involving photoperiod, temperature, autonomous floral initiation and gibberellins (Wilkie et al., 2008). However, photoperiodic induction has rarely been demonstrated in trees; most commonly, interactions between environmental stimuli (light intensity) and endogenous developmental cues (daily carbon assimilation) exert some control over floral initiation. Light intensity influences floral initiation in perennials, as described for apple, kiwifruit, olive and ornamental trees. In olive and ornamental trees, both high and low light intensity treatments reduced floral initiation (Stutte and Martin, 1986; Henriod et al., 2003). Similarly, in the black pepper cultivar *P. nigrum* cv. Reyin-1, the greatest flowering quantity occurred at intermediate light intensities (Zu et al., 2016). Although variations in light intensity can affect floral initiation intensity, they most likely are not the inductive stimuli in these cases; instead, a secondary factor, perhaps one related to assimilate production, stimulates flowering (Wilkie et al., 2008). Carbohydrates play roles in the regulation of plant metabolism. Increased amounts of stored carbohydrates can increase floral initiation, and light intensity has been reported to affect levels of stored carbohydrates in some horticultural trees (Tiessen and Padilla-Chacon, 2012; Hendriks *et al.*, 2003; Sato and Yanagisawa, 2014). However, it is unclear whether increased flowering intensity in adaptable light intensity treatments is due to the regulation of carbohydrates in black pepper.

Anthesis is one of the most important developmental events in the plant life cycle and involves the integration of environmental and developmental signals. Most plants are capable of utilizing the energy of light and storing it in carbon bonds through photosynthesis (MacNeill et al., 2017). Plants inventory available carbon resources prior to initiating the floral transition and will only commit to reproduction if there is enough stored energy within vegetative tissues (Yu *et al.*, 2000). Furthermore, despite having initiated the floral transition, plants can still alter carbon source-sink ratios during environmental changes depending on the developmental stage (MacNeill et al., 2017).

Two key components of carbohydrates are starch and sucrose. Starch metabolism is important in regulating key developmental processes, such as the floral transition. Leaf transitory starch is remobilized during reproductive development, and alterations in source starch reserves influence reproductive development (MacNeill et al., 2017). The sugar contents in the shoot apex have been correlated with flowering time, which is thought to occur through changes in starch remobilization or changes in photosynthetic carbon fixation (Lejeune et al., 1993; Bodson and Outlaw, 1985; Corbesier et al., 1998). Starchless leaves delayed flowering (Schlosser *et al.*, 2012), which illustrated the contribution of carbon remobilization from transitory starch to flowering (Braun *et al.*, 2014). To fully understand how carbohydrates regulate the floral transition, detailed physiological experiments are still necessary (Ohto et al., 2001). Sugar has been suggested to promote the floral transition in many plant species but to inhibit the floral transition in some plant species. This inhibition is probably not due to a suppression of photosynthetic activity (Jang et al., 1997); instead, it is likely due to growth caused by high concentrations of sucrose significantly delaying flowering time, causing an increase in the number of leaves at the time of flowering. Sucrose also affects a specific part of the vegetative phase, such as the transition from the adult vegetative phase to the reproductive phase. Further, the effects of sugar on the floral transition also differ depending on the concentration of sugar, the genotype of plants, and when in the vegetative growth phase the treatment is given (Ohto et al., 2001). Sucrose synthesis facilitates the rapid response of flowering to environmental conditions and its coordination with photosynthesis. However, little is known about the effects of shading on sucrose and starch metabolism in major source leaves of black pepper that affect the floral transition.

Carbohydrate metabolism enzymes have been studied extensively. Sucrose phosphate synthase (SPS) (EC 2.4.1.14) is a key regulatory enzyme involved in carbon partitioning between sucrose and starch in leaves (Huber and Huber, 1996). It catalyzes the penultimate step in sucrose synthesis and shares control of this pathway with the first committed step catalyzed by cytosolic FBPase (Liu et al., 2013). The crucial functions of sucrose synthase (SS) (EC 2.4.2.13) in plant metabolism are sucrose cleavage and energy provisioning (Li et al., 2002). SS contributes to maintaining sink strength and synthesizing storage products during organ maturation (Koch, 2004). Invertase (EC 3.2.1.26) hydrolyzes sucrose into hexoses. Invertase plays diverse roles in development and signaling (Ruan et al., 2010). All of these enzymes are affected by light, but the responses of the activities of these enzymes to light are different among plants and/or organs (Hubbard et al., 1989). To fully understand how the key enzymes in carbohydrate metabolism in major source leaves regulate sugar content and then control the floral transition in black pepper, detailed experiments are still necessary.

This study aimed (1) to study the effect of shading on the floral transition during the black pepper juvenile phase; (2) to clarify the relationship between sucrose metabolism and floral quantity under shading treatments; and (3) to identify key enzyme indicators that could be used regulate floral quantity in different developmental stages. These results might elucidate the physiological and biochemical mechanisms of major source leaves under shaded environments that could help in the regulation of inflorescence quantity by shading or other available measures.

## Results

### Effects of shading on pepper inflorescence number and length

Shading could significantly affect young pepper inflorescence and flower quantity during the annual growing period. In S1, the numbers of inflorescences were 42.81%, 86.82% and 91.30% lower under the intermediate and weak light conditions (30%, 60% and 75% shading intensities, respectively) than in the CK, while the length of inflorescences was significantly inhibited by 63.21% under the 75% shading intensity. In S2, the number of inflorescences was significantly inhibited, as shading intensity increased from 30% to 75%; specifically, the numbers were 49.62%, 91.54% and 88.38% lower than that in the CK, respectively. However, shading had little effect on inflorescence length in this period. In S3, the 30%, 60% and 75% shading intensities significantly reduced the number of inflorescences by 72.02%, 90.85% and 95.24%, respectively. The 60% and 75% shading intensities had significantly lower inflorescence lengths than did the CK (33.16% and 64.77% lower, respectively). In S4 and S5, a shading intensity lower than 30% led to very few alterations in the number and length of inflorescences. However, the number and length of inflorescences decreased by 99.16% and 100%, respectively, in S4 under the higher shading intensities (60% and 75%), while they declined by 80.67% and 93.66% in S5. Therefore, the optimum shading intensity based on conditions that significantly reduced the number of inflorescences was more than 30% in S1, S2, S3 and S4. Correspondingly, a shading intensity of more than 60% was appropriate for S5 (Fig. 1).

**Fig. 1.**
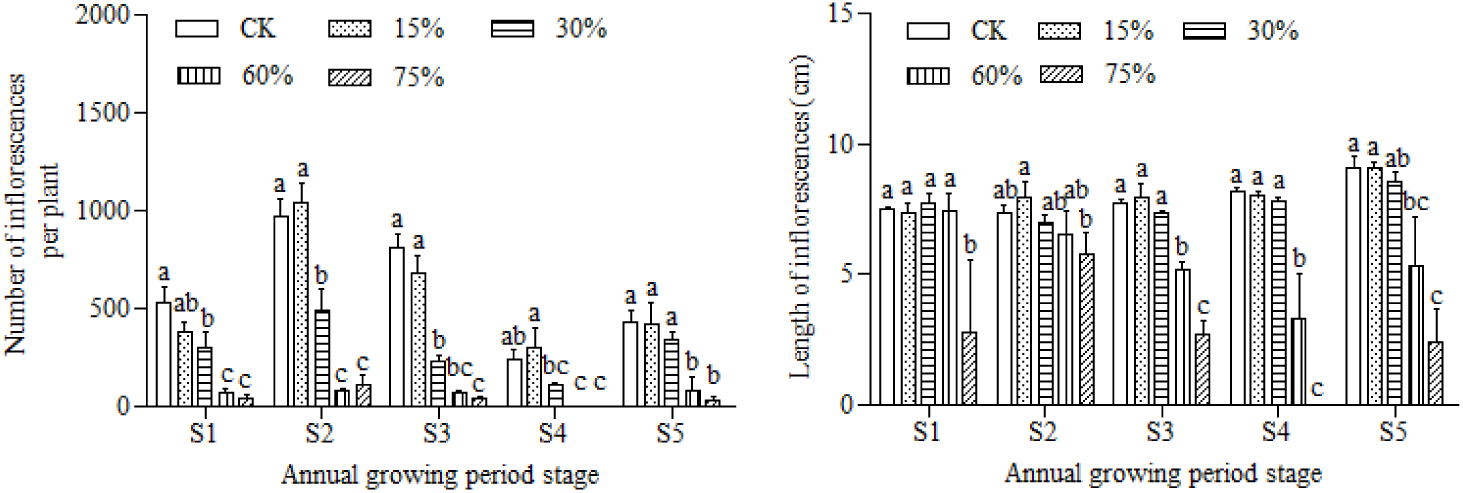
Effects of shading on black pepper inflorescence and flower quantity. Error bars indicate s.e.m. (n = 3); columns with different letters are significantly different according to ANOVA performed in SAS (*P* ≤ 0.05).

### Effects of shading on carbohydrate contents in major source leaves

The carbohydrate content changes in young pepper leaves under different shading intensities during the stages of the annual growing period are shown in Fig. 2. In S1, a decreasing trend in soluble sugar content with an average decreases of 19.26% and 15.54% occurred under the 60% and 75% shading intensities, respectively. Starch content also decreased under the higher shading intensities (60% and 75%), with reductions of 29.73% and 32.05%, respectively (Fig. 2c).

**Fig. 2.**
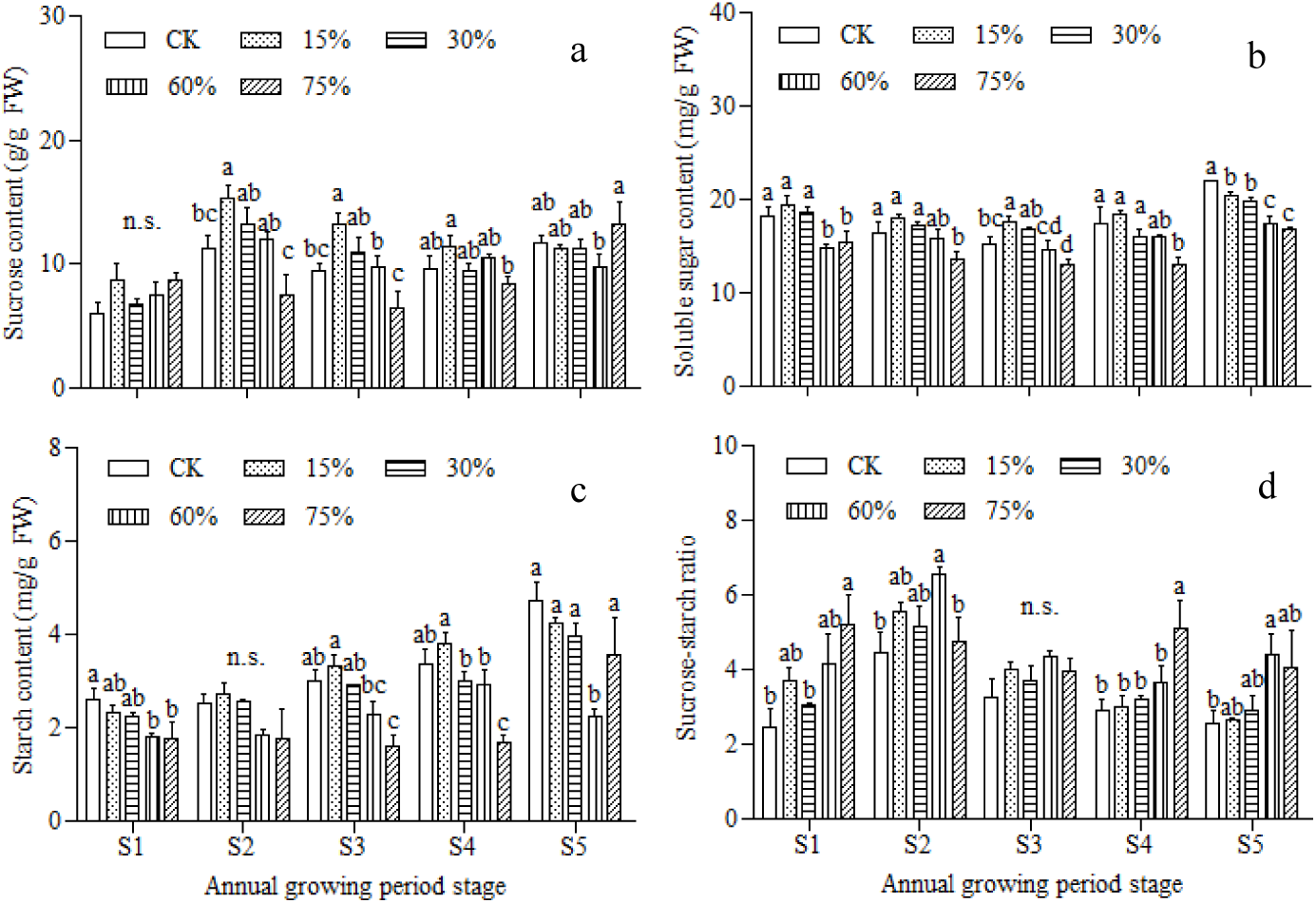
Effects of shading on carbohydrate contents in major source leaves of black pepper. Error bars indicate s.e.m. (n = 3); columns with different letters are significantly different according to ANOVA performed in SAS (*P* ≤ 0.05).

However, the sucrose-starch ratios were 52.87%, 24.59%, 70.90%, and 114.34% higher under the increasing shading intensities than in the natural sunlight CK (Fig. 2d). In S2, the sucrose content was 36.10% higher under the 15% shading intensity than in the CK (Fig. 2a). The soluble sugar content under the low shading intensity was similar to that in the CKbut was lower under the high shading intensity. The average value was 16.08% lower than that of the control. Shading had little effect on starch content. However, the sucrose-starch ratios were 25.39%, 15.73%, and 47.42% higher under the 15%, 30% and 60% shading intensities than in the CK (Fig. 2d). In S3, the sucrose content was 39.41% higher on average under the 15% shading intensity than in the CK, and the soluble sugar content was 14.87% higher on average under this shading intensity than in the CK. However, the soluble sugar content was 15.07% lower and the starch content was 46.15% lower under the 75% shading intensity than in the CK (Fig. 2b, 2c). In S4, there was no significant difference between lower shading treatments and the natural sunlight CK. The highest shading intensity of 75% inhibited the soluble sugar and starch contents, reducing them by 25.10% and 50.00%, respectively, compared with the control, but it increased the sucrose-starch ratio by 76.12% (Fig. 2b, 2c, 2d). In S5, the differences in sucrose content between the shading treatments and the CK were diminished, while the soluble sugar content under the shaded treatments was even lower than that in the CK. Specifically, the soluble sugar contents were 6.52%, 9.35%, 20.88% and 22.94% lower on average under the increasing shading intensities, respectively, than in the CK. The starch content decreased by 52.22% under the 60% shading intensity (Fig. 2c). The sucrose-starch ratio increased by 72.83% and 60.63% under the 60% and 75% shading intensities compared with that in the natural sunlight CK.

### Effects of shading on sucrose-metabolizing enzyme activities

Sucrose synthesis, partitioning and accumulation in black pepper leaves are influenced by the activities of four major sucrose-metabolizing enzymes. The results on the activities of these four enzymes are presented in the following sections. In S1, the SPS activity was 69.35% lower under the 75% shading intensity than in the CK. The SS activity decreased by 70.32% on average under the higher shading intensities (60% and 75%). There were no obvious effects of shading on NI and AI activities in this period. In S2, shading did not significantly reduce SPS, SS or NI. The intermediate shading intensity (30%) decreased AI activity by 31.48% in this period. In S3, the 30% and 60% shading intensities decreased SS activity by 56.46% on average. NI activity was reduced by 21.57%, 31.33% and 23.53% under the 30%, 60% and 75% shading intensities, respectively. However, the 75% shading intensity increased AI activity by 43.15% in this period. In S4, SPS was increased by 256.06% under the 75% shading intensity. AI activity declined by 44.91% on average under the 60% and 75% shading intensities. In S5, SS activity was decreased by 40.17% on average under the 60% and 75% shading intensities. The 30% shading intensity reduced AI activity by 36.55% (Fig. 3).

**Fig. 3.**
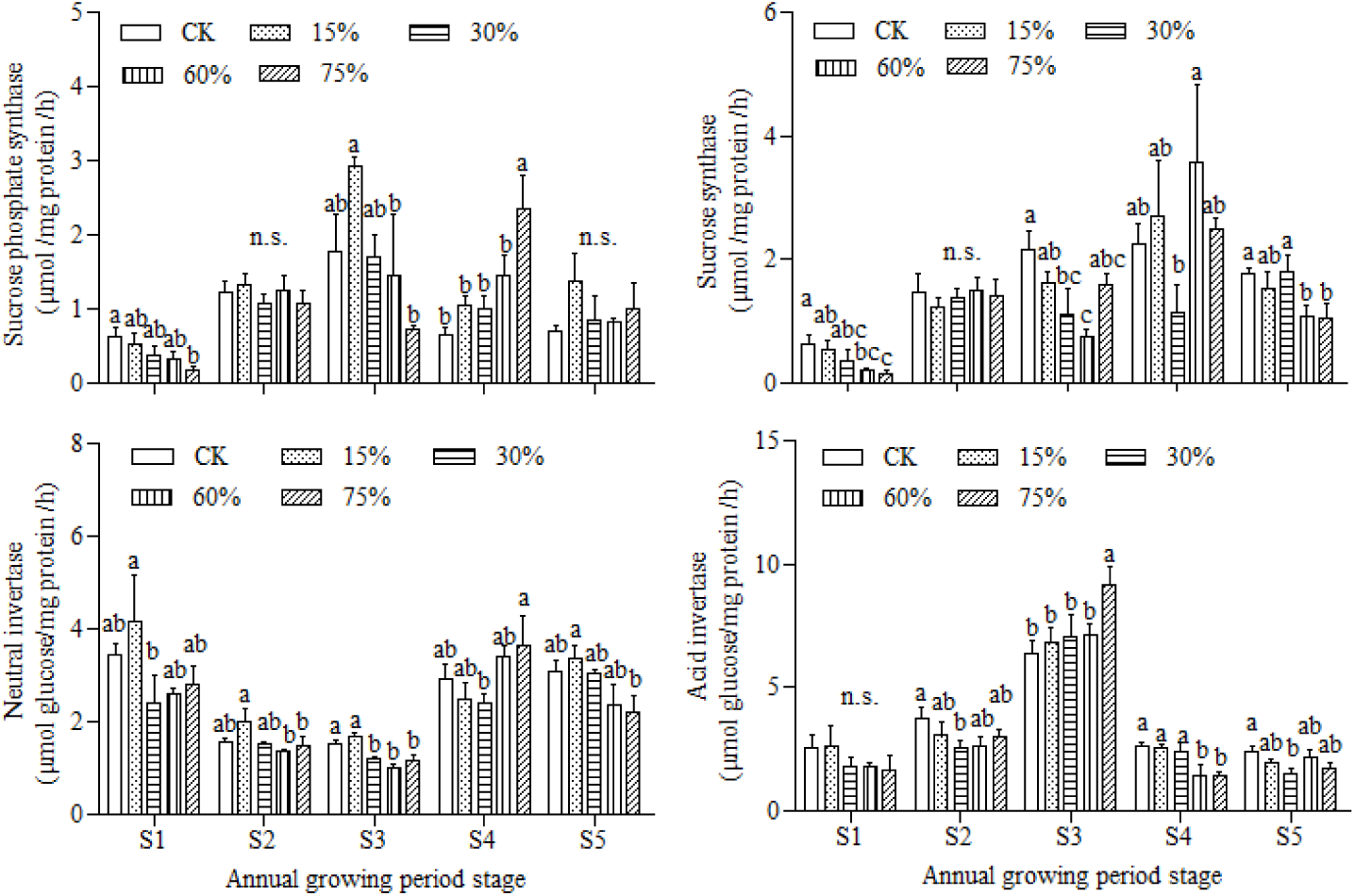
Effects of shading on sucrose-metabolizing enzyme activities in major source leaves of black pepper. Error bars indicate s.e.m. (n = 3); columns with different letters are significantly different according to ANOVA performed in SAS (*P* ≤ 0.05).

### Correlations between the activities of carbohydrate contents and floral quantity

The correlations between sucrose, soluble sugar, starch content and floral quantity were analyzed, and the results are listed in Table 1. In S1, significant positive correlations were observed between inflorescence number and soluble sugar and between inflorescence number and starch content. However, inflorescence length was only significantly and positively correlated with starch content. A significant negative correlation was observed between the floral quantity and the sucrose-starch ratio in S1, S4 and S5. Inflorescence number and inflorescence length were positively and significantly correlated with soluble sugar and starch content in S2, S3, S4 and S5. The inflorescence length was positively and significantly correlated with not only soluble sugar and starch content but also sucrose content in S3. The results showed that starch content was a key factor affecting floral quantity in S1, S2, S3 and S4. The sucrose-starch ratio affected floral quantity in S1, S4 and S5. Soluble sugar affected floral quantity in S5.

**Tab. 1.** Correlation coefficients of sucrose, soluble sugar and starch with floral quantity

| Annual growing period stage | Floral quantity | Sucrose | Soluble sugar | Starch | Sucrose-starch ratio |
| --- | --- | --- | --- | --- | --- |
| S1 | Inflorescence number | -0.376 | 0.662** | 0.778** | -0.716** |
|  | Inflorescence length | -0.239 | 0.459 | 0.684** | -0.759** |
| S2 | Inflorescence number | 0.473 | 0.571* | 0.635* | -0.313 |
|  | Inflorescence length | 0.439 | 0.542* | 0.678** | -0.341 |
| S3 | Inflorescence number | 0.433 | 0.514* | 0.720** | -0.431 |
|  | Inflorescence length | 0.726** | 0.806** | 0.927** | -0.295 |
| S4 | Inflorescence number | 0.234 | 0.530* | 0.642** | -0.562* |
|  | Inflorescence length | 0.352 | 0.658** | 0.834** | -0.809** |
| S5 | Inflorescence number | -0.130 | 0.789** | 0.608* | -0.627* |
|  | Inflorescence length | -0.306 | 0.738** | 0.556* | -0.665** |

Note: \* and \*\* indicate significance at the 0.05 and 0.01 levels, respectively.

### Relationship between carbohydrate content and sucrose-metabolizing enzyme activity

In S1, decreased SS activity could depress soluble sugar and starch content. The 60% and 75% shading intensities significantly decreased SS activity and then restrained inflorescence number and length. Therefore, the activity of SS can be controlled by shading to regulate floral quantity during this period. Decreased AI activity could obviously reduce soluble sugar and starch content, and shading had a nonsignificant effect on AI activity. Thus, shading is not the best way to limit AI activity, and we should look for other measures to control AI to regulate floral quantity (Tab.2). In S2, reduced AI activity was beneficial for reducing soluble sugar and starch content. The 30%, 60% and 75% shading intensities decreased AI activity and inflorescence number. Among these shading intensities, 30% shading significantly decreased the AI activity. Shading reduced soluble sugar and starch contents, resulting in a decrease in floral quantity. Therefore, regulating AI activity by shading is a suitable method to control floral quantity (Tab. 2).

In S3, decreased SPS activity decreased the soluble sugar and starch content. The 60% and 75% shading intensities decreased SPS activity and then reduced soluble sugar and starch content to control floral quantity. Therefore, the activity of SPS could be reduced by shading to regulate floral quantity during this period. Decreased NI activity could also reduce soluble sugar and starch content. The 30%, 60% and 75% shading intensities significantly reduced NI activity and floral quantity. Therefore, the decreased NI activity achieved via shading could be used to regulate floral quantity. AI activity was negatively correlated with soluble sugar and starch content and was increased under the 30% shading intensity and significantly improved under the 75% shading intensity. Therefore, improved AI activity under a 75% shading intensity could also reduce floral quantity in S3 (Tab. 2).

In S4, increased SPS activity and decreased AI activity decreased the soluble sugar and starch content but increased the sucrose-starch ratio. SPS activity was significantly improved, and AI was obviously inhibited by the 75% shading intensity. Therefore, a 75% shading intensity could lead to decreases in soluble sugar and starch content and an increase in the sucrose-starch ratio, thereby reducing floral quantity by increasing SPS activity and decreasing AI activity (Tab. 2).

In S5, improved NI activity could decrease sucrose content and the sucrose-starch ratio but increase soluble sugar and starch content. The 60% and 75% shading intensities inhibited NI activity. Among these shading intensities, the 60% shading intensity significantly decreased soluble sugar and starch content but increased the sucrose-starch ratio. In contrast, the 75% shading intensity obviously decreased the soluble sugar content. Inflorescence number and length decreased under the two shading intensities. Therefore, NI activity could be regulated to control floral quantity (Tab. 2).

### Regulation indicators and appropriate shading intensities in different growing periods

Based on an analysis of the correlation between the carbohydrate content of young pepper leaves and metabolic enzymes, combined with the effects of shade on leaf carbohydrate content, sucrose metabolism-related enzyme activity and floral quantity, indicators of sugar enzyme regulation and the appropriate shading intensity could be determined for different developmental stages in young black pepper.

In S1, the 60% and 75% shading intensities significantly decreased SS activity, but SS activity had no significant relationship with starch content. AI activity was not obviously inhibited under the two shading intensities, but AI activity was clearly related to soluble sugar and starch content, indicating that we should look for other methods to control AI activity. AI was the key indicator that could be used to control starch content and thus to suppress flowering. In S3, the 30% shading intensity significantly decreased NI activity to control flowering, while the 60% and 75% shading intensities inhibited SPS activity and improved AI activity but also decreased the vegetative tissue biomass of pepper plants (Fig. S1); therefore, other measures should be adopted to inhibit SPS activity and increase AI activity to control flowering (Tab. 3).

## Discussion

Floral induction responds to multiple external and endogenous signals to optimize the timing of the transition from vegetative to flower growth. The coordination of vegetative growth for carbohydrate accumulation and the timely transition to flower is critical for reproductive success. Floral inductive cues originate in mature leaves, and signals are transduced to the shoot apex to activate floral identity genes and initiate inflorescence development (Coneva et al., 2012). The carbohydrate levels in the shoot apex and mature leaves have been correlated with flowering time, which is thought to occur through changes in starch remobilization, photosynthetic carbon fixation (Lejeune et al., 1993; Bodson and Outlaw, 1985; Corbesier et al., 1998), or photosynthate partitioning (Rösti *et al.*, 2007). In this article, we added several novel findings about the effects of light on the regulation of floral quantity by carbohydrate activity in the major source leaves of black pepper (*P. nigrum* L.) in the juvenile phase.

Sucrose is the pivotal form of carbohydrate transported from photosynthetic sources (i.e., mainly mature leaves) to nonphotosynthetic sinks. Once carbohydrates have reached those sinks, sucrose must be degraded into hexoses to serve as metabolites to synthesize essential compounds including starch (Ruan et al., 2010; Ruan et al., 2014). Starch is also the major form of carbohydrate in plants and exists as either transitory or permanently stored starch (Streb and Zeeman, 2012). Transitory starch is a vital integrator of plant growth, buffering recurrent changes in carbon and energy availability that result from the diurnal light/dark rhythm (Geigenberger, 2011; Streb and Zeeman, 2012). In the chloroplasts of photosynthetic tissues, transitory starch accumulates gradually during the day and is consumed for continued sucrose biosynthesis and energy production at night (Hedhly et al., 2016; Thalmann et al., 2016). There are two enzymes in many plant species that catalyze the cleavage reaction of sucrose: SS and AI (Ruan et al., 2010). Our current understanding indicates that SS and AI are mainly involved in starch biosynthesis (Hedhly et al., 2016; McLaughlin and Boyer, 2004a; McLaughlin and Boyer, 2004b). In this work, we discovered that in S1 (the period dominated by vegetative growth), the 60% and 75% shading intensities markedly decreased floral quantity and the starch content in mature leaves and increased the sucrose content in leaves. There was a significant positive correlation between leaf starch and floral quantity. The sucrose-starch ratio in the mature major source leaves had an obvious inverse relationship with inflorescence quantity. A decrease in the SS activity in major source leaves could increase sucrose content and markedly decrease starch content, while a decrease in AI activity could slightly reduce sucrose content but significantly decrease starch content; therefore, reducing the activities of these two enzymes could increase the sucrose-starch ratio and markedly decrease starch content, thereby effectively controlling floral quantity. We determined that shading inhibits photosynthesis, which decreases SS and AI activities in the major source leaves, not only decreasing leaf starch content and increasing the sucrose-starch ratio but also further reducing inflorescence quantity (Fig. S2).

During early reproductive development, a plant may abort early reproductive tissue (pollen grains or ovules) and decrease sucrose transport to seeds to reduce sink strength and maternal carbon investment (Sun *et al.*, 2004). In S2 (the period dominated by anthesis), the 30% shading intensity significantly decreased floral quantity and AI activity without any reduction in vegetative growth. There was a significant positive correlation between leaf starch and floral quantity. Decreased AI activity could reduce starch content and then effectively control floral quantity. The results emphasize the importance of starch metabolism and source-sink relationships in regulating the key period dominated by anthesis (MacNeill et al., 2017).

Our research determined that an increased sucrose-starch ratio in the sources could not only delay flowering time but also reduce inflorescence quantity. However, sucrose can be maintained at a certain level in the sources (Park et al., 2009). In S3 (the period of fruit development), a high sucrose level inhibited AI activity under the 30% shading intensity. Therefore, AI could not catalyze starch synthesis. The upregulation of starch content in black pepper leaves might due to increased NI activity. To decrease starch content, we should decrease NI activity with a 30% shading intensity. In our research, we found a positive correlation between SPS activity and starch content in S3. SPS is a key regulator of the sucrose-starch ratio. However, in rice and tomato, reduced SPS activity resulted in higher starch accumulation (Coneva et al., 2012; Hashida et al., 2016). In hybrid poplar, increased SPS activity resulted in increased sucrose and starch accumulation in the leaf later in the growing season, which ultimately affected bud flush and senescence (Park et al., 2009). Our result is consistent with that of Park et al. To maintain the sucrose balance and decrease starch content, a 30% shading intensity should be adopted to reduce SPS activity.

In general, plants under stress during the grain-filling stage will experience increased source starch degradation in their vegetative tissues to be used to complete seed development (Cuellar-Ortiz *et al.*, 2008; Trouverie and Prioul, 2006). In this study, we aimed to inhibit flowering during this period. Therefore, the leaf starch content should be reduced. SPS has been shown to mediate the sucrose-starch ratio in plant leaves (Hashida et al., 2016). SPS can be both activated and deactivated by protein phosphorylation under osmotic stress and light, respectively (Huber and Huber., 1996). In S4 (the grain-filling stage), the 75% shading intensity could improve the SPS activity in major source leaves and, in turn, reduce starch content, increase sucrose content and the sucrose-starch ratio, and ultimately control floral quantity in black pepper. AI could promote sucrose utilization and starch biosynthesis (Ruan et al. 2010; McLaughlin and Boyer, 2004b). The 75% shading intensity could suppress the AI activity of major source leaves and in turn, leading to sucrose accumulation and starch utilization to control floral quantity.

NI is mainly involved in starch biosynthesis and sucrose decomposition (McLaughlin and Boyer, 2004b; Hedhly et al., 2016). Rearrangements of starch metabolism are observed when plants are subjected to short periods of osmotic stress. Soluble sugars may function as osmoprotectants during stress responses (Thalmann et al., 2016). In S5 (the fruit maturation stage), the 60% and 75% shading intensities could suppress the NI activity of major source leaves to decrease soluble sugar content, osmotic stress and starch content, thus control ling inflorescence length and quantity. Furthermore, the NI activity reduction could also increase the sucrose-starch ratio, allowing the control of floral quantity.

In conclusion, we found that in black pepper, shading results in starchless leaves, which leads to significant reductions in the number of flowers, illustrating the contribution of carbon remobilization from transitory starch to control floral quantity (Braun et al., 2014). Starch is the major component of black pepper (Zhu et al., 2018). Total starch content amounts to over 50% of dry black pepper drupes (Zhu et al., 2017). However, our current knowledge of the mechanisms controlling starch metabolism to control black pepper flowering is limited (Thalmann and Santelia, 2017). To fully understand how carbohydrates regulate the floral transition, detailed physiological experiments are still necessary (Ohto et al., 2001).

## Acknowledgments

This work was supported by the Natural Science Foundation of China (grant no. 31601820) and Central Public-interest Scientific Institution Basal Research Fund for Chinese

Academy of Tropical Agricultural Sciences (grant no.1630142017013).

## Supplementary information

### Effects of shading on vegetative tissue biomass accumulation in S3

In S3, compared with the control, the 15% and 30% shading intensities had no significant effect on vegetative tissue biomass accumulation; however, the 60% and 75% shading intensities significantly suppressed pepper plant growth (Fig. S1).

### Effect of shading on photosynthetic rate

Shading inhibited the photosynthetic rate in S1, S2 and S3. However, in S4 and S5, the photosynthetic rate was not suppressed under shading conditions (Fig. S2).

---

**Tab. 3.** Regulation indicators and appropriate shading intensity for the annual growing period stages

| Annual growing period<br>stage | Regulation indicator | Appropriate shading intensity |
| --- | --- | --- |
| S1 | SS and AI activity reduction | 60-75% |
| S2 | AI activity reduction | 30% |
| S3 | SPS and NI activity reduction & AI activity improvement | 30% |
| S4 | SPS activity improvement & AI activity reduction | 75% |
| S5 | NI activity reduction | 60-75% |

**Tab. 2.**
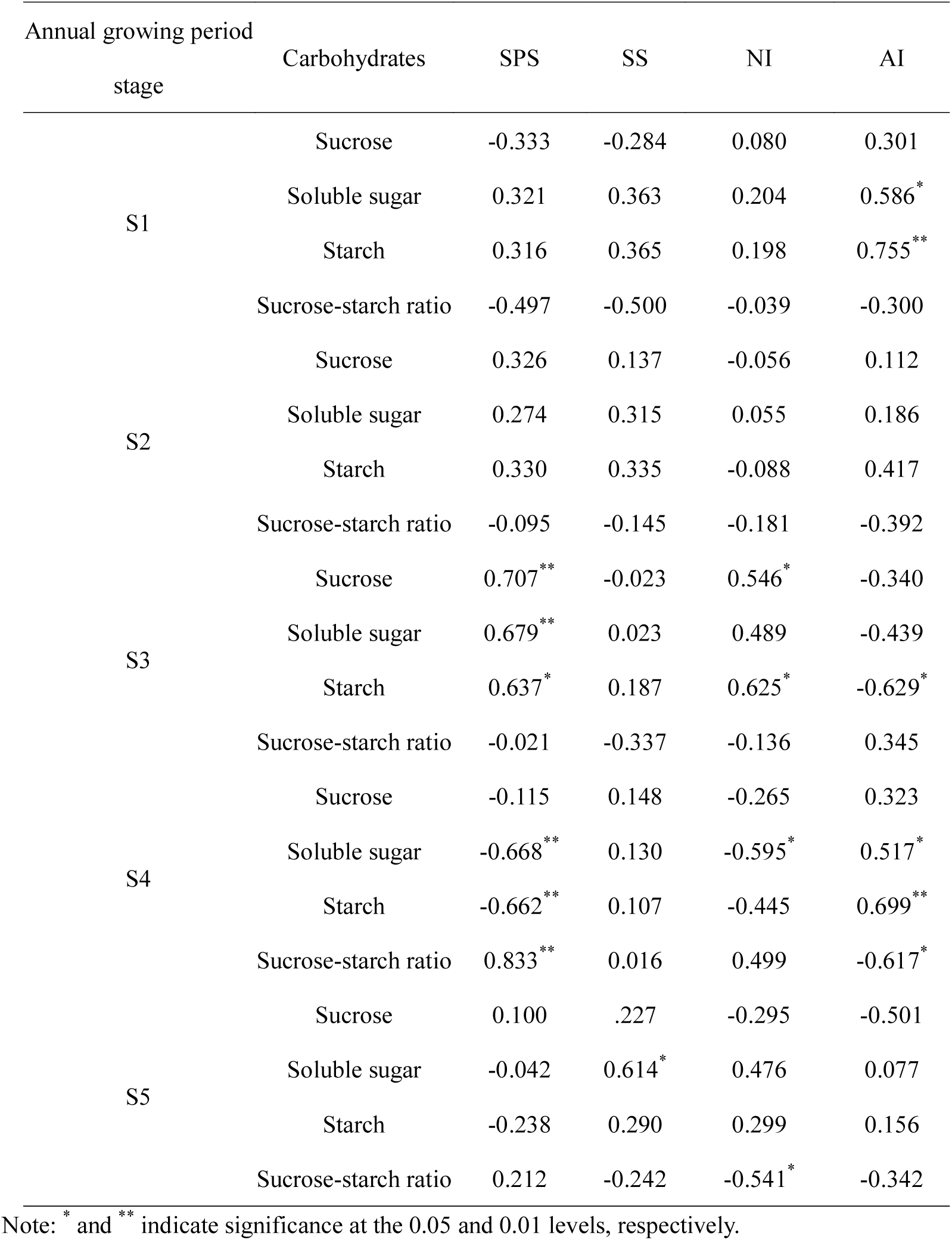
Relationship between carbohydrate content and sucrose-metabolizing enzyme activity

| Annual growing period<br>stage | Carbohydrates | SPS | SS | NI | AI |
| --- | --- | --- | --- | --- | --- |
| S1 | Sucrose | -0.333 | -0.284 | 0.080 | 0.301 |
|  | Soluble sugar | 0.321 | 0.363 | 0.204 | 0.586* |
|  | Starch | 0.316 | 0.365 | 0.198 | 0.755** |
|  | Sucrose-starch ratio | -0.497 | -0.500 | -0.039 | -0.300 |
| S2 | Sucrose | 0.326 | 0.137 | -0.056 | 0.112 |
|  | Soluble sugar | 0.274 | 0.315 | 0.055 | 0.186 |
|  | Starch | 0.330 | 0.335 | -0.088 | 0.417 |
|  | Sucrose-starch ratio | -0.095 | -0.145 | -0.181 | -0.392 |
| S3 | Sucrose | 0.707** | -0.023 | 0.546* | -0.340 |
|  | Soluble sugar | 0.679** | 0.023 | 0.489 | -0.439 |
|  | Starch | 0.637* | 0.187 | 0.625* | -0.629* |
|  | Sucrose-starch ratio | -0.021 | -0.337 | -0.136 | 0.345 |
| S4 | Sucrose | -0.115 | 0.148 | -0.265 | 0.323 |
|  | Soluble sugar | -0.668** | 0.130 | -0.595* | 0.517* |
|  | Starch | -0.662** | 0.107 | -0.445 | 0.699** |
|  | Sucrose-starch ratio | 0.833** | 0.016 | 0.499 | -0.617* |
| S5 | Sucrose | 0.100 | .227 | -0.295 | -0.501 |
|  | Soluble sugar | -0.042 | 0.614* | 0.476 | 0.077 |
|  | Starch | -0.238 | 0.290 | 0.299 | 0.156 |
|  | Sucrose-starch ratio | 0.212 | -0.242 | -0.541* | -0.342 |

Note: \* and \*\* indicate significance at the 0.05 and 0.01 levels, respectively.

